# Sociality, diurnal temperature range and isothermality: Significant determinants of mass-independent resting metabolic rate in subterranean African mole-rats (Superfamily Bathyergidae)

**DOI:** 10.64898/2026.02.13.705831

**Authors:** Jack E. Thirkell, Monica A. Daley, Nigel C. Bennett, Daniel W. Hart, Chris G. Faulkes, Steven J. Portugal

## Abstract

Animals exhibit a diverse range of sociality from the strictly solitary to the highly social. Different forms of sociality have evolved in response to ecological constraints and selective habitat pressures, which are governed by the energetic and fitness costs to an individual. Uniquely among mammals, the clade of African mole-rats (Bathyergidae and Heterocephalidae) covers three distinct life-history forms of sociality: solitary, social and eusocial species. This variety in social structure makes them a model clade to study how metabolic traits vary between different forms of sociality. Resting metabolic rates (RMR) of seven African mole-rat species, ranging from solitary to eusocial, were measured using open-flow respirometry. Results were combined with published data, enabling the inclusion and statistical analysis of 16 species in total. We identified distinct allometric scaling of RMR, with eusocial species exhibiting a considerably greater rise in RMR with increases in body mass. This is likely attributable to reproductive and behavioural divisions of labour, and mass-dependent colony roles in eusocial species. Phylogenetically-informed analyses further identified that sociality, in addition to select bioclimatic traits – diurnal temperature range (°C) and isothermality (%) – significantly explain variation in the mass-independent RMR of African mole-rats. These findings elucidate, for the first time, that sociality can be a determinant of RMR, and calls for further study to identify the wider significance of sociality on mammalian metabolism, as well as exploring the allometric scaling of metabolic rate with respect to mammalian sociality.

## Introduction

The adaptive benefits of group living are well-documented and include increased predator detection, higher offspring survival, greater efficiency of resource acquisition, enhanced thermoregulation and combinations thereof (Kotze *et al*. 2008; Sichilima *et al*. 2008; Meldrum & Ruckstuhl 2009). However, sociality does not occur without costs, as animal aggregations can lead to increased predation attempts, intraspecific resource competition and disease transmission (Alexander 1974), particularly under high-population densities (Dobson & Hudson 1995; Begon *et al*. 2002). Despite there being, under certain circumstances, energetic savings associated with being social, ultimately, the benefits of sociality must outweigh any energetic and fitness costs.

African mole-rats (Bathyergidae and Heterocephalidae) are subterranean rodents endemic to sub-Saharan and Eastern Africa (Faulkes *et al*. 1997). Although the Heterocephalidae diverged from the Bathyergidae approximately 31.2 million years ago, collectively they are a monophyletic clade (Superfamily Bathyergidae) that exhibit different forms of social living, as well as many shared behavioural and physiological traits (Bennett & Faulkes 2000; Patterson & Upham 2014; Rodrigues *et al*. 2015). While a subterranean environment offers respite from environmental extremes and enhanced protection from predation (Bennett & Faulkes 2000), it presents a unique set of challenges. Subterranean environments are characterised by the intermittence of resources, an absence of light, high humidity, reduced gas ventilation, low atmospheric oxygen and the increased energetic costs associated with moving through resistive substrates (Burda *et al*. 2007; Jonz *et al*. 2016; Merchant *et al*. 2024a). In response to the increased energetic demands, subterranean rodents tend to have significantly reduced mass-specific resting metabolic rates (msRMR; 0.4-1.2 ml O_2_ g^-1^.hr^-1^), compared with terrestrial rodents of a comparable size (1.0-1.4 ml O_2_ g^-1^.hr^-1^) (Bennett & Faulkes 2000; Kingma *et al*. 2012). The *respiratory stress hypothesis* (Arieli 1979), *cost of burrowing hypothesis* (Vleck 1979; Lovegrove 1987; Lovegrove & Wissel 1988) and *thermal stress hypothesis* (McNab 1966) are three hypotheses that have been proposed to explain why subterranean rodents have low RMRs. In addition, African mole-rats have lower body temperatures (T_b_), relative to most placental mammals, and higher thermal conductances (°C) than rodents of an equivalent body size (Šumbera 2019). These factors likely protect against heat stress in closed burrow systems (McNab 1966). While aspects of their fossorial-related physiology have been studied extensively at the individual species level, a holistic approach to determining the extent to which bioclimatic traits may be determining metabolic rate in this group is currently lacking.

As a group, African mole-rats are a unique mammalian clade that encompasses species with a wide spectrum of sociality, ranging from solitary to social and eusocial species (Bennett & Faulkes 2000). *Bathyergus*, *Georychus* and *Heliophobius* are solitary, *Cryptomys* and *Fukomys* are social, with the exception of *Fukomys damarensis*, which like the monotypic species of the genus *Heterocephalus* (*Heterocephalus glaber*) is eusocial (Faulkes *et al*. 1997). Given that molecular phylogenies have identified *F. damarensis* and *H. glaber* as being evolutionarily divergent (Allard & Honeycuttt 1992; Faulkes *et al*. 1997), eusociality is considered to have most likely evolved independently in these two species. Eusocial species can be defined by three characteristics: 1) cooperative brood care; 2) reproductive division of labour (often determined by body mass) and 3) overlapping generations (Michener 1969; Jarvis 1981), while more recently a eusociality continuum has been proposed as a more realistic approach of quantifying sociality across vertebrates and invertebrates (Sherman *et al*. 1995). Similarly, social species can be defined by group living, but exhibit a less strict dominance hierarchy and a loose to absent reproductive division of labour (Moolman *et al*. 1998; Faulkes & Bennett 2001). In solitary species, co-occupancy of the burrows occurs only briefly during the breeding season or when the females have young (Bennett & Jarvis 1988a). A further distinction between species of these three forms of sociality is their lifetime reproductive success - the proportion of animals that attain reproductive status, which ranges between 1-15% for eusocial species (1% for *H. glaber* and 15% for *F. damarensis*) (Jarvis & Bennett 1993; Jarvis *et al*. 1994), 40-60% for social species and approaches 100% for solitary species (N. C. Bennett, unpublished). This variety of social organisation makes African mole-rats an ideal group to study the metabolic traits associated with different forms of mammalian sociality.

A possible reason for the variation in RMR between different forms of sociality may be attributable to relative brain size. It has been proposed that encephalisation (increasing brain size relative to body mass), while variable across taxonomic groups, is associated with mammalian sociality (Shultz & Dunbar 2010). It is inferred that species exhibiting high sociality have greater metabolic rates, owing in-part to the energetics demands of an increased brain size. However, African mole-rats are an apparent exception to this pattern; sociality does not appear to drive the evolution of large brains in eusocial mole-rats (Kverková *et al*. 2018). Instead, the wide range of sociality observed among the genera of African mole-rats is considered an evolutionary response to different ecological constraints and selective habitat pressures (Bennett & Faulkes 2000). More specifically, the Aridity Food Distribution Hypothesis (AFDH) proposes that sociality in African mole-rats evolved in environments of low and unpredictable rainfall, where geophytes become irregularly distributed and the risks of unsuccessful foraging are high (Jarvis *et al*. 1994, 1998). Consequently, the energetic costs of foraging are greater than in areas of high rainfall; cooperative foraging in social and eusocial species is energetically advantageous under these environmental constraints (for alternative hypotheses, see Burda et al. 2000). Therefore, it could be considered that bathyergids would not follow the typical patterns established in mammals, whereby social species have comparatively larger brain sizes, and the associated energetic costs this demands.

Here we test the hypothesis that differences in metabolic rate arise between species with different forms of sociality, and identify whether the relationship between RMR and body mass varies between eusocial, social and solitary species. In eusocial species, we expect a steeper scaling exponent, attributable to reproductive and behavioural divisions of labour, and an allometric assignment of colony roles that is unique to eusocial species (Bennett & Jarvis 1988b; Scantlebury *et al*. 2006b; Faulkes & Bennett 2013). As neither social nor solitary species exhibit these divisions of labour, the relationship between RMR and body mass between these two groups is expected to be comparable. Finally, we investigate whether sociality and pertinent bioclimatic traits are significant determinants of metabolic rate in African mole-rats within a phylogenetically-controlled framework.

## Material and Methods

### (a) Study species (7-species)

The resting metabolic rates (RMR) of seven African mole-rat species: *Fukomys damarensis, Heterocephalus glaber, Cryptomys hottentotus hottentotus, Cryptomys h. mahali, Cryptomys h. pretoriae, Bathyergus suillus* and *Georychus capensis* were assessed. For this study we adopted categorical definitions of sociality (i.e. eusocial, social and solitary) for mole-rat species (Jarvis 1981; Jarvis & Bennett 1993), rather than a continuous measure (Sherman *et al*. 1995). Collectively, these species represent five of the six genera within the African mole-rat clade and encompass all forms of sociality.

All animals had been maintained in captivity for a minimum of one year at the time of the study, to ensure metabolic acclimation to captive conditions (Bennett *et al*. 1992, 1993b). With the exception of *H. glaber* all species were housed in large containers at the Department of Zoology and Entomology, University of Pretoria, where ambient laboratory temperature (T_a_) was maintained between approximately 22-25°C. *Heterocephalus glaber* were housed in burrow systems and at the School of Biological and Chemical Sciences, Queen Mary University of London, where laboratory T_a_ was maintained at approximately 30°C. Social and eusocial species were maintained in their respective colonies, while all species were provisioned with appropriate nesting material and *ad libitum* access to food.

### (b) Respirometry experimental procedure

Resting metabolic rate was determined through the measurement of the rate of oxygen consumption (V̇O_2_) and carbon dioxide production (V̇CO_2_), using an open-flow respirometer (Sable Systems International, Las Vegas, NV). Animals were fasted for >3hrs prior to RMR assessments, to ensure a post-absorptive state and exclude the potential influence of digestion on metabolic activity (Šumbera 2019). The rate of oxygen consumption was measured at ambient temperatures within each species respective thermoneutral zone (TNZ; Supplementary Table 1). Despite an apparent absence of circadian rhythms among these species (Bennett & Faulkes 2000), for continuity with other metabolic studies on African mole-rats and to follow established protocols, we conducted all assessments between 08:00 – 18:00hrs, to mitigate against the potential effects of endogenous metabolic rhythms. An absence of circadian rhythms of metabolism, along with the unknown extent of stress and variation in the time that animals were fasted, resulted in RMR, rather than basal metabolic rate (BMR), being a more applicable measure of energy expenditure in this study; the strict criteria for BMR could not be guaranteed (Šumbera 2019). Resting metabolic rate is considered to be between 10-25% greater than BMR, typically (Sherwood *et al*. 2005).

Respirometry chambers (repurposed airtight containers) measured 11.5L for *B. suillus*, 6.5L for *G. capensis* and *F. damarensis*, 2L for *C. h. hottentotus*, *C. h. mahali* and *C. h. pretoriae* and 0.9L for *H. glaber* (Thirkell *et al*. 2025; Merchant *et al*. 2024b). The different sized chambers catered for interspecies mass differences (Table 2 for mean body masses). Each chamber was fitted with 4-mm inlet and outlet ports. The outside air was pulled through the chambers at varying flow rates depending on the chamber size; 1,250 ml min^-1^ for *B. suillus*, 1,000 ml min^-1^ for *G. capensis* and *F. damarensis*, 750 ml min^-1^ for *C. h. hottentotus*, *C. h. mahali* and *C. h. pretoriae* and 600 ml min^-1^ for *H. glaber*.

Each respirometry assessment lasted approximately 65 minutes and consisted of a 10-minute baseline to assess ambient O_2_ level, a 45-minute metabolic assessment, followed by a further 10-minute baseline to reassess ambient O_2_. The analogue outputs of O_2_ (%), CO_2_ (%), flow rate (ml min^-1^), relative humidity (%), barometric pressure (kPa) and temperature (°C) were recorded concurrently using a universal interface (UI2, Sable Systems International, Las Vegas, NV). These measurements were sampled (1 Hz) and monitored in real-time using ExpeData software (Sable Systems International, Las Vegas, NV), which enabled the progress and stability of each animal’s respirometry trace to be visually assessed. Additionally, this enabled the manual addition of markers on the trace to note times of aberrant behavioural observations or external confounding factors. This real-time monitoring also safeguarded against potentially dangerous spikes in CO_2_ or drops in O_2_, at which point the assessment would have been terminated. Body mass (g) was measured immediately preceding each assessment using Oertling electronic weigh scales.

Incurrent airflow was controlled using a flow regulating pump (SS-4, Sable Systems International, Las Vegas, NV), calibrated against a certified mass flow meter (FoxBox, Sable Systems International, Las Vegas, NV), placed downstream of the respirometry chamber. Fractional concentration of O_2_ was measured using an oxygen analyser (FC-10a, Sable Systems International, Las Vegas, NV), which was calibrated to ambient air O_2_ concentration (20.95%) before each trial. Fractional concentration of CO_2_ was measured using a carbon dioxide analyser (CA-10a, Sable Systems International, Las Vegas, NV), and relative humidity measured using a water vapour analyser (RH-300, Sable Systems International, Las Vegas, NV). Barometric pressure and temperature were measured from inbuilt sensors in the FC-10a oxygen analyser. Anhydrous Indicating Drierite™ was used to scrub atmospheric water from the excurrent air between the water vapour and CO_2_ analysers, and again between the CO_2_ scrubber and the oxygen analyser (W. A. Hammond Drierite Company LTD, U.S.A). CO_2_ was scrubbed from the excurrent air between the CO_2_ and O_2_ analysers (Soda Lime, Sigma Aldrich, Merck KGaA, Darmstadt, Germany).

Data, once exported from ExpeData, were processed in Matlab (version 9.6. Natick, Massachusetts: The MathWorks Inc., 2019). O_2_ and CO_2_ were corrected for baseline drift and any time lag between these two variables (due to the delay in airflow between analysers) was corrected using cross-correlation. The fractional O_2_ signal was corrected for the removal of CO_2_ (O_2__corrected), the fractional CO_2_ signal was corrected for the removal of water vapour (CO_2__corrected), and the flow rate was corrected to Standard Temperature and Pressure (STP) conditions. A 5-minute minimum analysis region was selected for RMR, corresponding to the lowest stable O_2_ consumption and CO_2_ production, during which the animal was considered to be most restful. The average over this period was used to obtain RMR estimates (V̇O_2_ and V̇CO_2_), calculated using the formulae;

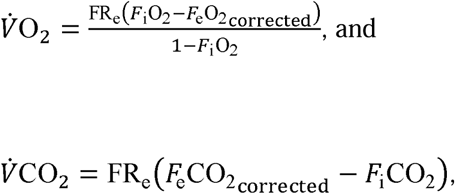

where *F*_1_ and *F*_e_ are incurrent and excurrent fractional concentrations (%) of O_2_ and CO_2_ (Lighton 2008). The ratio of V̇CO_2_ to V̇O_2_ determined the respiratory quotient (RQ) (Lighton 2008).

For the purposes of data presentation mean RMR values were calculated from individual measurements of V̇O_2_ and are, unless otherwise stated, presented as the mean ± SD, corrected to STP conditions.

### (c) Combined study (16-species)

Data from the 7-species assessment in the present study were combined with published RMR and body mass data from a total of 16 African mole-rat species. Where there were multiple studies of any one species, the mean RMR and body mass was calculated, weighted by the sample size of the respective studies. In most studies, RMR was referenced in ml O_2_ hr^-1^, however, there were two studies in which metabolic rates were referenced in either KJ hr^-1^ or KJ day^-1^ and therefore, an equivalent of 20.1 KJ L^-1^ O_2_ was used to convert energy values to oxygen consumption (Zelová *et al*. 2010, 2011).

### (d) Bioclimatic traits and multicollinearity

Geographic range distributions in the form of shapefiles (.shp) were sourced from the International Union for the Conservation of Nature (IUCN, 2019). There were two exceptions, *Heliophobious emini* and *Cryptomys hottentotus nimrodi*, for which no official population census had been undertaken. Instead, for these species, longitude and latitude coordinates were identified from published studies using populations of wild-caught animals ((Bennett *et al*. 1996; Ngalameno *et al*. 2017; Katandukila 2020)). Geographic coordinates from a total of five populations of *H. emini* and three populations of *C. h. nimrodi* were identified (Table 1).

**Table 1.** The geographic coordinates of wild-caught populations of *Cryptomys hottentotus nimrodi* and *Heliophobius emini*.

| Species | South | East | Latitude | Longitude | Reference |
| --- | --- | --- | --- | --- | --- |
| *C. h. nimrodi* | 20°17' | 28°56' | -20.283333 | 28.93333 | (Bennett *et al.* 1996) |
| *C. h. nimrodi* | 20°11' | 28°37' | -20.183333 | 28.61667 | (Bennett *et al.* 1996) |
| *C. h. nimrodi* | 22°10' | 29°30' | -22.166667 | 29.5 | (Bennett *et al.* 1996) |
| *H. emini* | 6°57'16.45" | 37°32'5.5" | -6.954569 | 37.53483 | (Katandukila 2020) |
| *H. emini* | 6°49'13.296" | 37°40'14.016" | -6.82036 | 37.67056 | (Ngalameno *et al.* 2017) |
| *H. emini* | 6°58'42.888" | 37°33'0.036" | -6.97858 | 37.55001 | (Ngalameno *et al.* 2017) |
| *H. emini* | 6°58'48.108" | 37°32'46.859" | -6.98003 | 37.54636 | (Ngalameno *et al.* 2017) |
| *H. emini* | 6°58'48.18" | 37°32'46.859" | -6.98005 | 37.54635 | (Ngalameno *et al.* 2017) |

A total of 32 pertinent bioclimatic traits were initially considered for inclusion (Supplementary Table 1). These were sourced and calculated from high resolution spatiotemporal climate data: WorldClim Global Climate Data 1970-2000 (Fick & Hijmans 2017), ERA5-Land 1950-2021 (Muñoz Sabater 2019, 2021) and Socioeconomic Data and Application Center (Imhoff *et al*. 2004; Imhoff & Bounoua 2006). While a broad range of bioclimatic traits were included, these were restricted to only those that had either been documented previously to have had a significant effect on mammalian RMR, or could be reasoned to have an effect in these subterranean study species: annual temperature, diurnal temperature range, isothermality, temperature seasonality, maximum temperature, minimum temperature, annual temperature range, temperature of wettest, driest, warmest and coldest quarter, annual precipitation, precipitation of wettest and driest month, precipitation seasonality, precipitation of wettest, driest, warmest and coldest quarter, altitude, net primary productivity (NPP), temperature 2m above surface, skin reservoir content (SRC), skin temperature (i.e. surface temperature), soil temperature and volumetric soil water at 0-7cm, 7-28cm, 28-100cm and 100-289cm depths. For each bioclimatic trait a median across each species’ geographic range distribution was calculated, or in the case of *C. h. nimrodi* and *H. emini*, the mean at each population’s latitudinal and longitudinal coordinates. For species that had two or more distinct geographic range distributions or populations, a mean was then used to calculate a species average (Supplementary Table 2).

Multicollinearity between bioclimatic traits was assessed using a variation inflation factor (VIF) in the R package *car* (v. 3.0-7) (Fox & Weisberg 2019), in a two-step process. Given the relatively small number of mole-rat species assessed within this study (N=16), bioclimatic traits were first broadly grouped into four categories: I) temperature, II) precipitation, III) subterranean environment, and IV) miscellaneous traits (Supplementary Table 1). The VIF assessed collinearity within each group, where traits with the largest VIF >5 were removed in a stepwise process, until only traits with a VIF <5 remained. These remaining bioclimatic traits from each of the four groups were then combined and the VIF of this new group similarly assessed. Twenty-five of the initial traits were determined to be collinear; seven bioclimatic traits (maximum temperature, precipitation of the driest and coldest quarter, volumetric soil water (0-7cm) skin reservoir content, diurnal temperature range and isothermality; Supplementary Table 3) were retained in PGLS modelling, along with sociality. An unavoidable drawback of our study is that there are only two species of eusocial mole rats. We ran a standard leverage analysis to determine if the eusocial species, or indeed any species, was largely influencing the overall relationship. The hat values (all <2 - the standard threshold) suggest that none of the 14 mole-rat species had high leverage.

### (e) Statistical analyses and phylogenetic method

All calculations and statistical analyses were performed in R statistical software (v. 3.5.21) (RStudio 2020).

#### Metabolic allometry

To account for the considerable range of body masses across the 16 species (37.5-695.9g), and the non-normal distribution of both RMR and body mass values, these weighted mean values were logarithmically transformed (log_10_). To assess differences in the allometric scaling of metabolic rates between eusocial, social and solitary species, separate linear regressions of RMR against body mass were constructed for each of the three social classes and their scaling exponent identified.

#### Phylogenetic tree

An ultrametric phylogenetic tree was constructed for the 16 species assessed within this study using the Open Tree of Life, using the R package *rotl* (Michonneau *et al*. 2016) (Figure 1). The Open Tree of Life synthesises published phylogenies along with taxonomic data.

**Figure 1.**
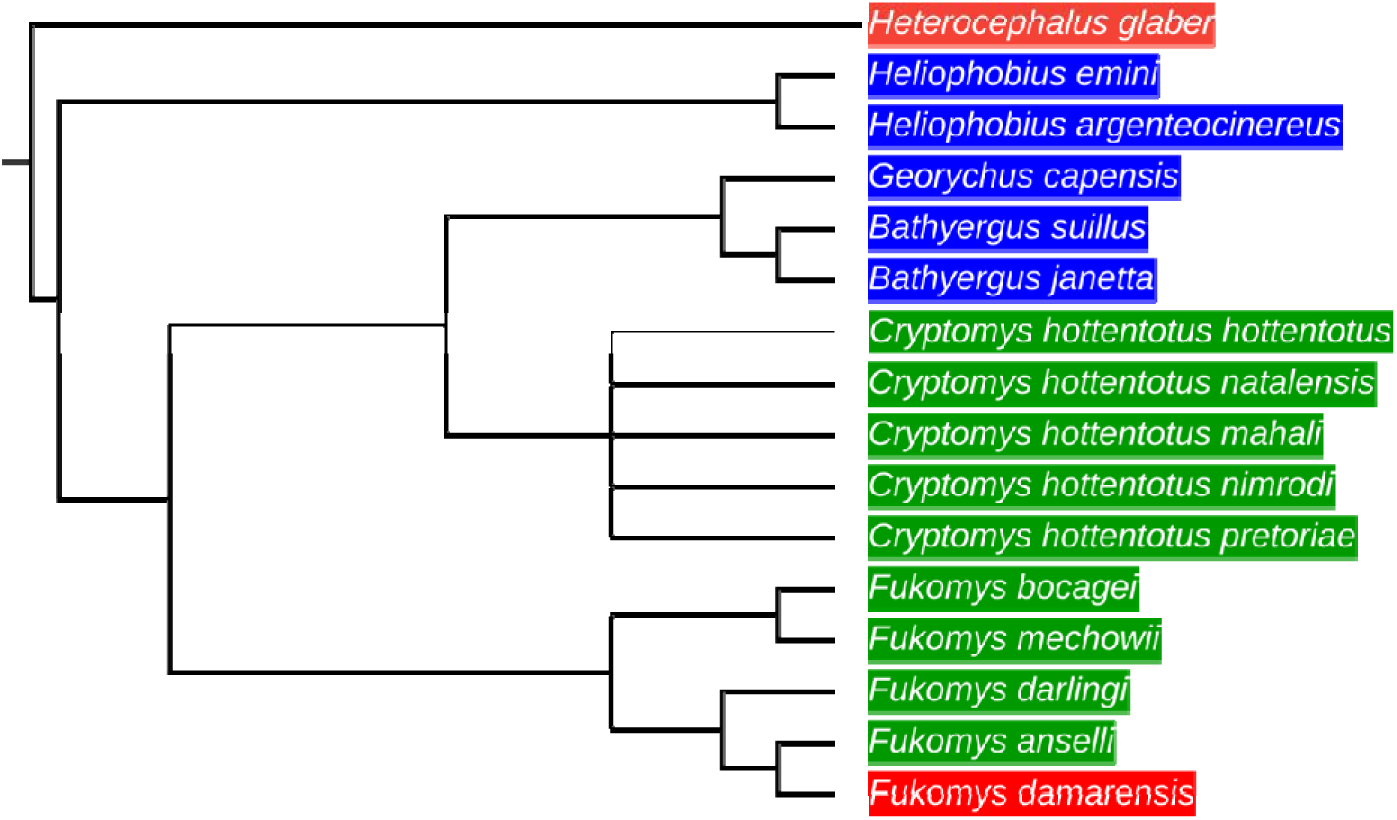
An ultrametric phylogeny of 16 African mole-rat species (Bathyergidae and Heterocephalidae) assessed within this study. Phylogenetic tree was constructed using the Open Tree of Life (see Material and Methods). Tip labels highlighted in red denotes eusocial species, green denotes social species and blue denotes solitary species.

#### Phylogenetic generalised least squares (PGLS)

Despite body mass being the single greatest determinant of mammalian metabolic rate, this study set out to identify to what extent bioclimatic traits describe the remaining variation in metabolic rate. Body mass was accounted for in the PGLS modelling by regressing RMR against body mass, and calculating the residual for each species (i.e. mass-independent RMR); mass-independent RMR was used as the response trait in all PGLS models.

Phylogenetic generalised least squares (PGLS) modelling was performed using the R package *caper* (v. 1.0.1) (Orme *et al*. 2018). To assess the phylogenetic signal of mass-independent RMR across the 16 species, Pagel’s lambda (λ) was calculated using the R package *phytools* (v. 0.6-60) (Revell 2012) on 999 phylogenetic simulations. Pagel’s λ is an index (0 ≤ λ ≥ 1) used to determine the strength of the phylogenetic signal of a trait across a phylogeny. Pagel’s λ values approaching 0 indicates no phylogenetic signal (i.e. mass-independent RMR has evolved independently of phylogeny), whereas, Pagel’s λ values approaching 1 indicates the evolution of a trait according to Brownian motion (Molina-Venegas & Rodríguez 2017).

Phylogenetic generalised least squares models were used to check the explanatory power of different combinations of up to three predictor traits for variation in mass-independent RMR (n = 92; Supplementary Table 4 for full list of PGLS models). The explanatory power of the statistical models were ranked based on Akaike Information Criterion corrected for small sample sizes (AICc) (Akaike 1973). Furthermore, a conditional average model was constructed that incorporated all models with a δAICc less than or equal to two. Model selection and averaging was performed using the R package *MuMIn* (v. 1.43.6) (Bartoń 2019).

## Results

### (a) Metabolic assessment of seven-species

The resting metabolic rate (RMR; ml O_2_ hr^-1^) of 118 animals across seven species of African mole-rat was measured (Table 2; see Supplementary Figure 1 for graphical representation of the intra- and interspecific distribution of RMR and body masses).

**Table 2.** The mean resting metabolic rate (RMR; ml O_2_ hr^-1^) and body mass values across seven African mole-rat species.

| Species | Sample size | Sociality | RMR (ml O<sub>2</sub> hr<sup>-1</sup>) | Body mass (g) |
| --- | --- | --- | --- | --- |
| *B. suillus* | 13 | Solitary | 467.67 ± 150.66 | 668.46 ± 212.81 |
| *C. h. hottentotus* | 12 | Social | 139.31 ± 50.5 | 77.42 ± 10.88 |
| *C. h. mahali* | 10 | Social | 112.44 ± 32.32 | 92.9 ± 18.94 |
| *C. h. pretoriae* | 15 | Social | 100.88 ± 17.73 | 103.33 ± 25.12 |
| *F. damarensis* | 27 | Eusocial | 257.25 ± 79.56 | 124.26 ± 32.23 |
| *G. capensis* | 20 | Solitary | 212.33 ± 51.03 | 150.5 ± 53.24 |
| *H. glaber* | 21 | Eusocial | 45.71 ± 17.54 | 31.33 ± 9.69 |

### (b) Sociality and allometric analysis of resting metabolic rate

A total of 49 published mean RMR and body mass values were identified, in 35 respective studies of 16 species. These were combined with the mean RMR and body mass values of the seven species assessed in this study (Table 3 for summary details of contributing studies. See Supplementary Table 5 for full details of contributing studies).

**Table 3.**
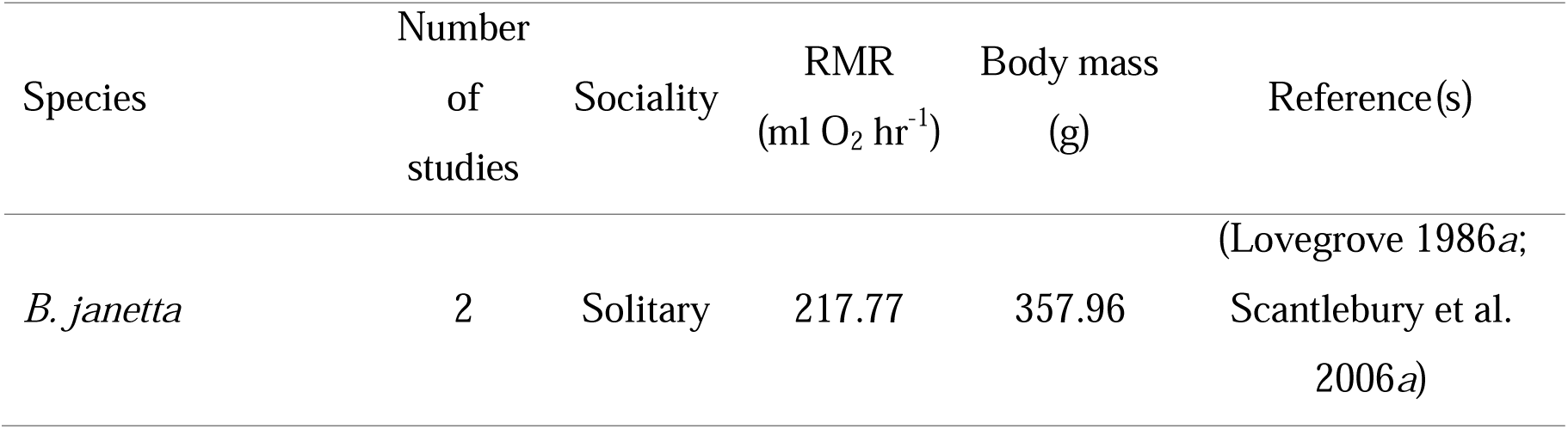
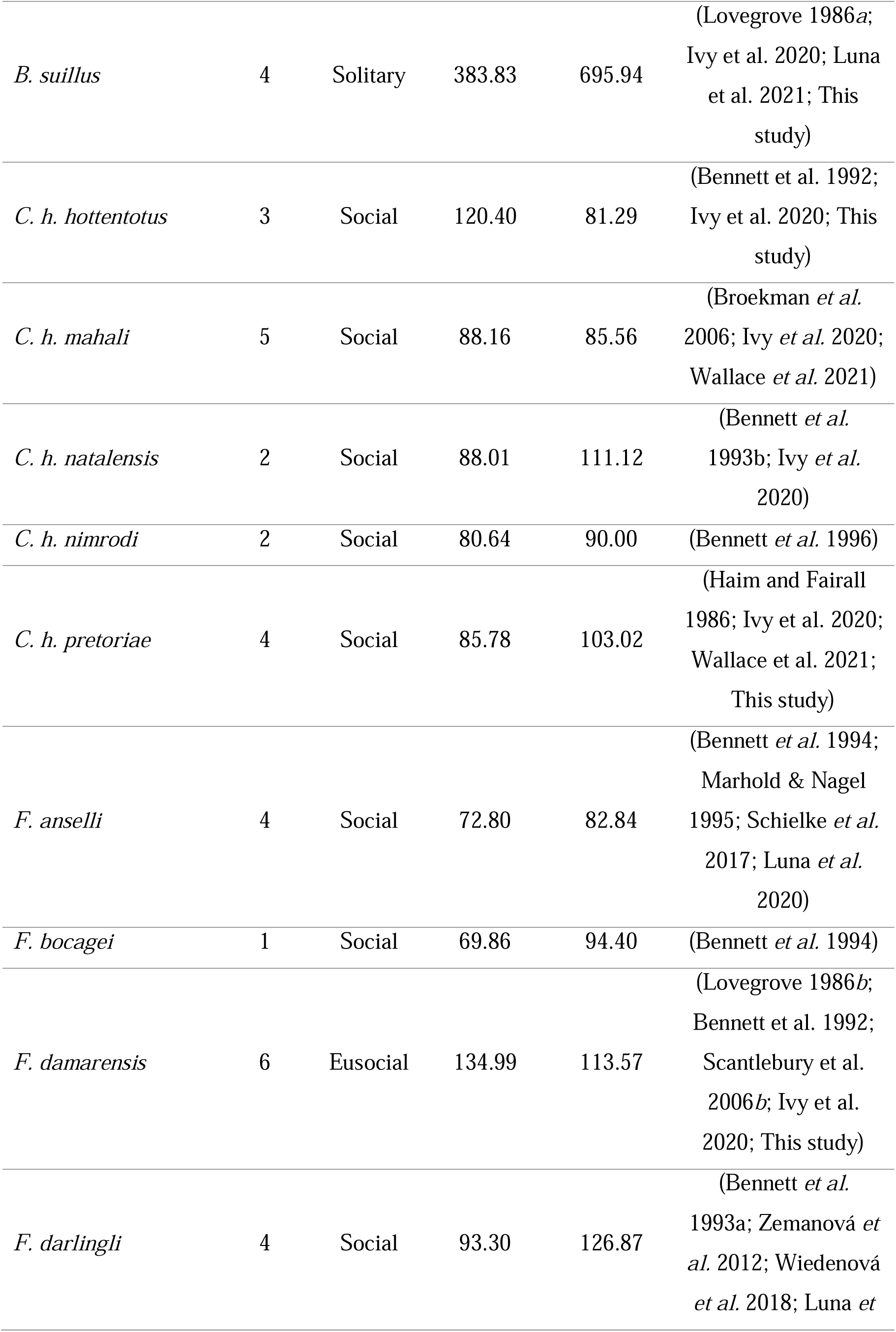
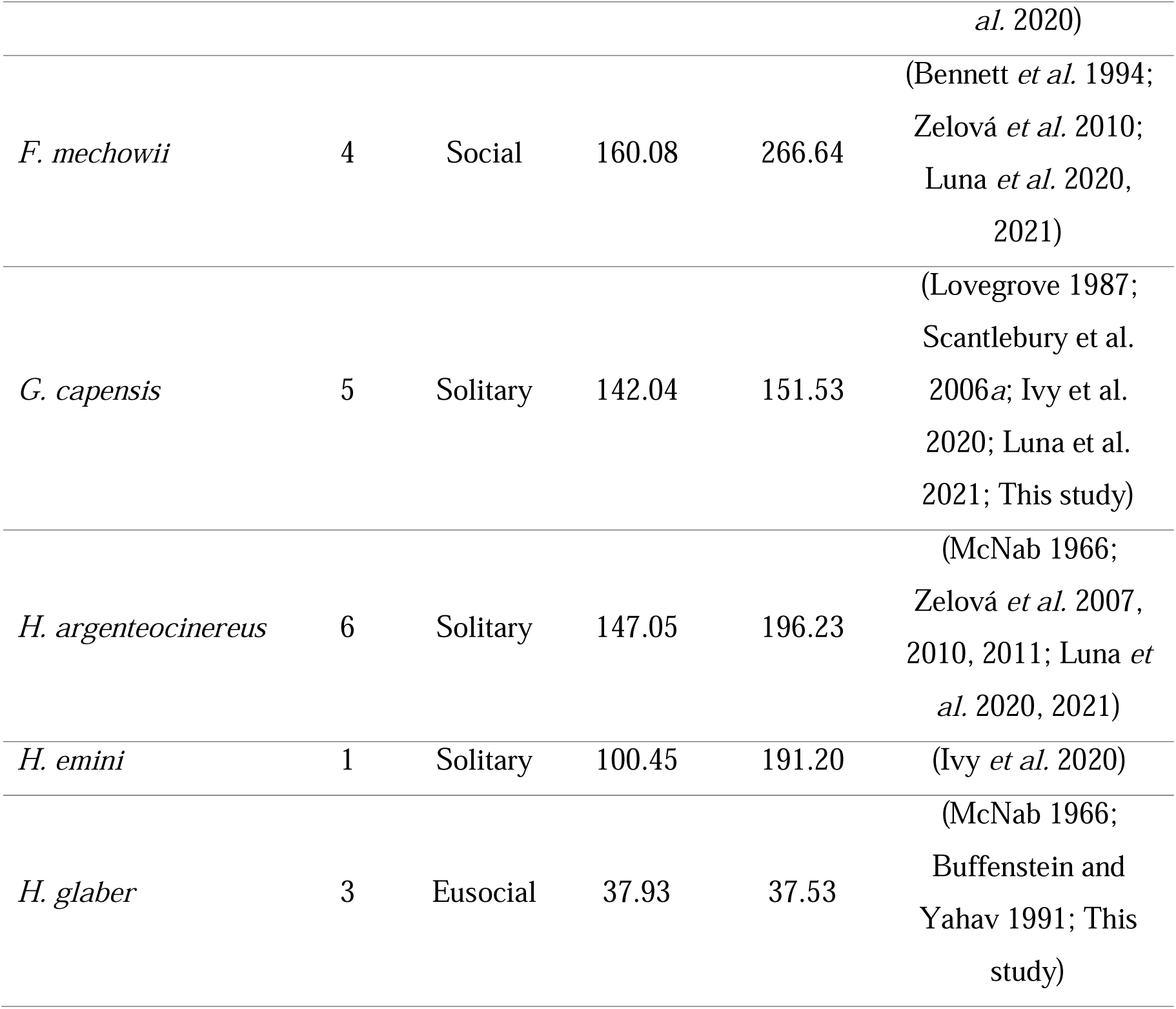
The mean metabolic and body mass values of 16 African mole-rat species, weighted by the sample size of the respective studies.

When RMR was regressed against body mass the resultant scaling exponent was greatest in eusocial species (1.15 ml O_2_ g^-1^.hr^-1^; Figure 2), indeed, allometric scaling of metabolic rate was twice as steep as that of social species (0.51 ml O_2_ g^-1^.hr^-1^; Figure 2). Whereas, the scaling exponent of solitary species was 0.77 ml O_2_ g^-1^.hr^-1^ (Figure 2). This demonstrates that eusocial species exhibit a much greater rate of increase in their RMR for a given increase in body mass, compared with social and solitary species.

**Figure 2.**
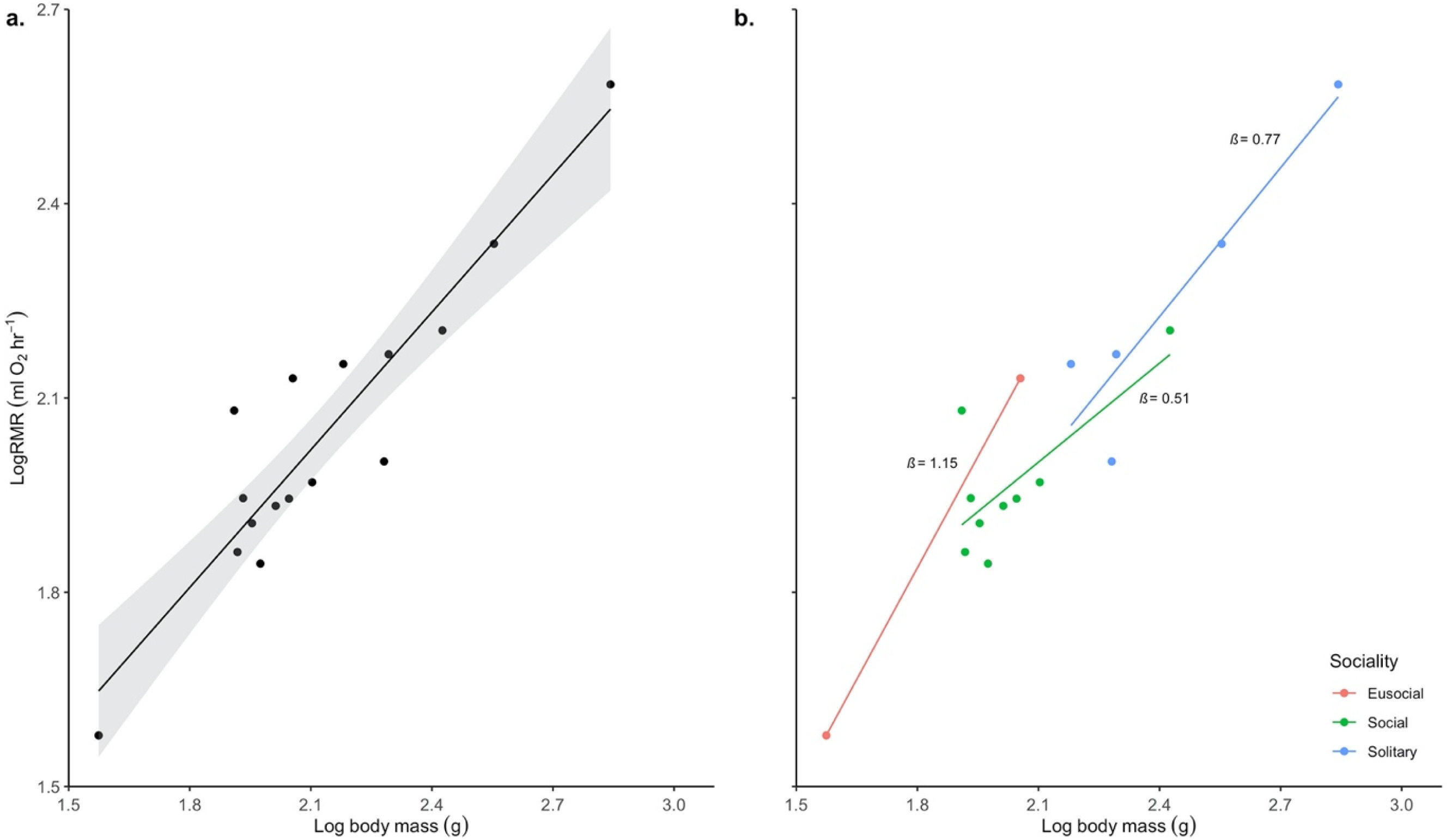
Regressions of resting metabolic rate (RMR) against body mass in a) 16 African mole-rat species, b) stratified by sociality, where solitary species are denoted in blue (*N* = 5), social species in green (*N* = 9) and eusocial species in red (*N* = 2). Metabolic rate (ml O_2_ hr^-1^) and body mass (g) values were logarithmically transformed (log10).

### (c) Phylogenetic generalised least squares (PGLS) models

Pagel’s λ for mass-independent RMR was calculated at less than 0.001, indicating that phylogeny (Figure 1) had no significant effect on mass-independent RMR. Five top-ranking PGLS models (δAICc ≤ 2) were identified (Table 4), suggesting that the mass-independent RMR of African mole-rats is predicted by isothermality, sociality, diurnal temperature range, skin reservoir content and volumetric soil water. However, model averaging subsequently indicated that only isothermality (*z* = 2.45, p = 0.01; Table 5), sociality [social – eusocial] (*z* = 2.75, p = 0.01; Table 5) and diurnal temperature range (*z* = 2.10, p = 0.04; Table 5) were significant determinants of mass-independent RMR within this mammalian clade.

**Table 4.** Top-ranked phylogenetic generalised least squares (PGLS) models of mass-independent resting metabolic rate (δAICc ≤ 2), which contributed to the conditional average model (Table 5). The relative model weights (W) are estimates across the entire set of 92 PGLS models (Supplementary Table 5).

| Model | DF | loglik | AICc | Delta AICc | Model weight |
| --- | --- | --- | --- | --- | --- |
| Isothermality | 4 | -67.28 | 146.2 | 0 | 0.16 |
| Isothermality + Sociality | 6 | -63.13 | 147.6 | 1.38 | 0.08 |
| Diurnal temperature range | 4 | -68.24 | 148.1 | 1.91 | 0.06 |
| Isothermality + Skin reservoir content | 5 | -66.07 | 148.1 | 1.94 | 0.06 |
| Isothermality + Volumetric soil water | 5 | -66.08 | 148.2 | 1.96 | 0.06 |

**Table 5.** While skin reservoir content and volumetric soil water were retained in the top-ranked models (Table 4), model averaging indicated that neither of these traits were significant determinants of mass-specific resting metabolic rate in 16 African mole-rat species (Bathyergidae and Heterocephalidae).

|  | Estimate | Std. Error | z value | Pr(>\|z\|) |
| --- | --- | --- | --- | --- |
| (Intercept) | 114.90 | 106.30 | 1.08 | 0.28 |
| Isothermality | -2.36 | 0.96 | 2.45 | 0.01 |
| Sociality [Social vs Eusocial] | -35.08 | 12.75 | 2.75 | 0.01 |
| Sociality [Solitary vs Eusocial] | -20.49 | 12.52 | 1.64 | 0.10 |
| Sociality [Solitary vs Social] | 14.59 | 8.48 | 1.72 | 0.09 |
| Diurnal temperature range | 6.98 | 3.32 | 2.10 | 0.04 |
| Skin reservoir content | -184200 | 126200 | 1.46 | 0.14 |
| Volumetric soil water | -92.07 | 63.45 | 1.45 | 0.15 |

## Discussion

This is the first study to demonstrate differences in the allometric scaling of metabolic rates associated with social or group living and paves the way for further study on such metabolic traits in other animal systems. Specifically, the relationship between resting metabolic rate (RMR; ml O_2_ hr^-1^) and body mass exhibited by eusocial species is distinct from solitary and social species. That is to say, eusocial species have a considerably greater scaling exponent (1.15 ml O_2_ g^-1^.hr^-1^), compared to solitary (0.77 ml O_2_ g^-1^.hr^-1^) or social species (0.51 ml O_2_ g^-1^.hr^-1^). It is possible that the observed differences are attributable to the specific thermal biology of naked mole-rats, or other physiological factors, rather than eusociality alone. Indeed, the two eusocial species are morphologically dissimilar; *F. damarensis* is considerably heavier than *H. glaber* (Table 2) and is entirely covered in fur. While *H. glaber* has a similarly higher thermal conductance to other social Bathyergidae species, a recent review concluded that *H. glaber* is not as energetically distinct as previously presumed, instead it is the absence of fur and small body mass that distinguishes this species from others (Šumbera 2019).

With this in mind, we posit that the higher scaling exponent associated with eusociality is likely related to reproductive and behavioural divisions of labour, and an allometric assignment of colony roles, whereby an individual’s role within the colony varies with body mass (Bennett & Jarvis 1988b; Scantlebury *et al*. 2006b; Faulkes & Bennett 2013). This interpretation is supported by a recent study that showed a complex relationship between body size and work-related behaviours in naked mole-rats, in which the probability of an individual being observed working was predicted by both age and body mass-cubed, with larger individuals less likely to be observed working (Gilbert *et al*. 2020); furthermore, high-ranking individuals worked for shorter durations. A reduction in colony work would infer a concurrent reduction in activity-related energy expenditure, which could be redirected towards the morphological, physiological and behavioural preparation associated with breeding, dominance challenges and dispersal (O’Riain *et al*. 1996). Previous work also established that the predominant role of larger individuals shifts from burrow maintenance to colony defence or dispersal (O’Riain & Jarvis 1997; Faulkes & Bennett 2013; Mooney *et al*. 2015), thus these individuals are likely to need a higher metabolic capacity to effectively exhibit these behaviours. Furthermore, we speculate that body compositional changes are likely associated with changing colony role, not just total mass or size gains. Defence and dispersal behaviours likely require an increase in both body mass and relative musculature, which would be more metabolically demanding to maintain at rest, as well as energetically costly to transport. The allometric assignment of colony roles may necessitate a large increase in RMR across the species’ relatively small range in body mass. Although this study assessed metabolic rates at rest, future research would benefit from identifying whether there are interspecies differences in both the net-cost of transport and body composition, in addition to whether there is discernable metabolic variation between the different roles in a colony exhibited by eusocial species.

Our *a priori* prediction was that the relationship between body mass and RMR would be similar in social and solitary species, on the basis that neither group exhibits reproductive or behavioural divisions of labour. However, we identified that solitary species exhibit a greater relative increase in RMR for a given increase in body mass (Figure 2). The apparent difference between solitary and social species could be governed by environmental constraints. Whereas social species more often occur in environments of greater aridity, solitary species tend to occur in mesic and temperate environments that are characterised by increased variability in both daily and seasonal temperature (Bennett *et al*. 1988; Lovegrove & Knight-Eloff 1988; Lovegrove & Wissel 1988; Roper *et al*. 2001; Šumbera *et al*. 2004). It is this seasonal variability in burrow temperature that has resulted in the marked breeding seasonality exhibited by solitary species (Bennett & Faulkes 2000; Sumbera *et al*. 2003). Despite burrow systems being substantially buffered against variable above-ground temperatures (Gates 1963; Bennett & Faulkes 2000; Roper *et al*. 2001), burrow temperatures of solitary species tend to get colder during an austral winter compared to that of social and eusocial species (Šumbera 2019), which could necessitate morphological, physiological and behavioural adaptations (Merchant *et al*. 2024c; Merchant *et al*. 2024d). In the absence of social thermoregulation, solitary species have developed alternate thermoregulatory strategies; solitary species, including, *B. suillus*, *G. capensis* and *Heliophobius argenteocinereus*, have a denser and longer pelage to reduce heat exchange (Bennett *et al*. 2006, 2009; Šumbera 2019), tend to have an increased body mass and have a wider range of TNZs (Bennett & Faulkes 2000; Šumbera 2019). Ultimately, differences in metabolic traits between solitary and social species are likely to relate to variation in life-history traits, driven by ecological constraints and selective habitat pressures.

Phylogenetic generalised least squares modelling, which accounted for body mass, indicated that phylogeny is not a significant determinant of RMR in African mole-rats (λ < 0.001). This is perhaps intuitive given that eusociality is believed to have arisen twice within Bathyergidae and Heterocephalidae (Allard & Honeycuttt 1992; Faulkes *et al*. 1997). While three other bioclimatic traits were included in the modelling: maximum temperature (°C) and precipitation (mm) in the driest and coldest quarters, such traits did not improve the fit of the models (δAICc ≥2). We speculate that differences in the composition and insulation-type qualities of the substrates in which the different species inhabit, in addition to intraspecific thermoregulatory mechanisms (i.e. fur densities, tunnel and burrow depths, huddling, etc.), may be attributing to the relative non-significance of these traits. Although skin reservoir content (m of water equivalent) and volumetric soil water (0 – 7cm; m^3^ m^-3^) were retained in the top-ranked PGLS models, model averaging indicated that neither of these traits were significant determinants of mass-independent RMR (Table 5). However, it was determined that sociality, isothermality and diurnal temperature range were traits that significantly explained variation in mass-independent RMR in African mole-rat species.

Even despite the relatively conserved range of body masses across this group of species (35g – 1,600g), there are observable differences in allometric scaling. We recognise that the inclusion of only two eusocial species is a limitation and that only tentative interpretations can be made of the metabolic scaling exponent of eusociality. However, as *F. damarensis* and *H. glaber* are the only two recognised eusocial mammals, increasing the statistical power of the study by increasing the number of eusocial species assessed was not possible. As all animals had been maintained in captivity for a minimum of one year prior to metabolic assessment, it is worth noting possible effects of captivity on RMR. Indeed, *Cryptomys h. hottentotus* and *C. h. natalensis* have been shown to have a reduced mass-specific RMR (msRMR) following a period of acclimation to captivity (Bennett *et al*. 1992, 1993b), while conversely, *F. damarensis* have been documented to have an increased msRMR (Bennett *et al*. 1992). Nevertheless, it is hoped that this study lays the foundations for future studies to explore the effects of sociality in other mammalian clades; with the addition of further species, across a wider range of body masses, clearer differences in the allometric scaling of metabolic rate could become evident between groups of species with different social classes. Notwithstanding, phylogenetically-informed analysis suggest that sociality is a significant determinant of mass-independent metabolic rate; a species’ sociality affects its metabolic rate, at least amongst African mole-rats.

## Conclusion

This assessment of seven species describes, to date, the most extensive metabolic study of African mole-rats, accounting for two solitary, three social and two eusocial species; an unavoidable drawback of our study is that there are only two species of eusocial mole rats (and indeed, mammals) for comparison. Despite the metabolic costs of many life-history traits having already been studied, the energetics associated, specifically, with sociality has not previously been investigated. We demonstrate, for the first time, the possible existence of distinct allometric scaling of metabolic rates between different forms of sociality. We conclude that the higher scaling exponent between RMR and body mass, exhibited in eusocial species, is likely attributable to reproductive and behavioural divisions of labour, and an allometric assignment of colony roles, which are absent in both social and solitary species. Furthermore, through phylogenetically-informed modelling, sociality, isothermality and diurnal temperature range were identified to be significant determinants of mass-independent RMR in African mole-rats. It is suggested that future studies should explore whether similar observations between different forms of sociality are observed across the mammalian kingdom.

## Ethics

Experimental procedures involving live animals and data collection described herein were approved by Royal Holloway University of London, Queen Mary University of London and the University of Pretoria Animal Ethics Committee (Ref. EC004-19). The study was conducted in accordance with appropriate institutional and national guidelines.

## Data accessibility

All code and data supporting this article is included as electronic supplementary material.

## Authors’ contributions

J.E.T. collected data, conducted statistical analyses and drafted the manuscript. S.J.P. conceptualised the study, contributed to study design and critically reviewed the final draft. M.A.D, C.G.F, N.C.B and D.W.H contributed to study design and facilitated data collection through the provision of respirometry equipment, a MATLAB script and study animals. Interpretation and preparation of the manuscript was conducted by J.E.T and supported by all authors. All authors have contributed significantly to the study, have reviewed the manuscript and have agreed to its submission.

## Acknowledgements

We would like to show our gratitude to Dr Rudiger Riesch, Dr Sarah Papworth and the Portugal Lab Group for useful discussion, and the BSU staff at Queen Mary and technical staff at the University of Pretoria for care of the animals. Special thanks also to Dr Stephanie McClelland for assistance with the phylogenetic modelling and to Eleanor Dixon for valued contributions to data collection.

