## Supplementary Figure 1 for "Sociality, diurnal temperature range and isothermality: Significant determinants of mass-independent resting metabolic rate in subterranean African mole-rats (Superfamily Bathyergidae)"

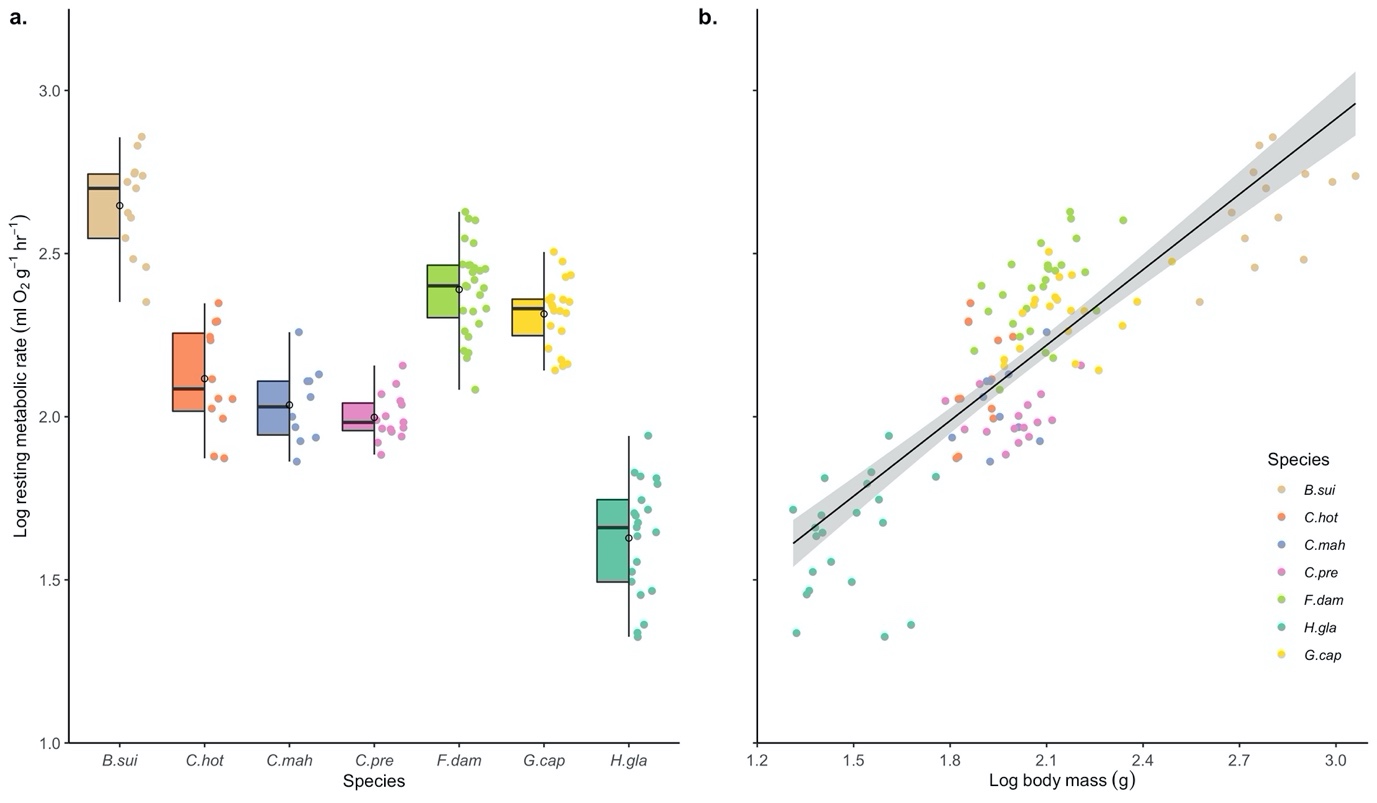

Supplementary Figure 1. The distribution of resting metabolic rate and body mass values across seven African mole-rat species, represented by a a) jitterplot and b) scatterplot. Metabolic and body mass values were logarithmically transformed (log10), and mean values are indicated by open circles. *B. sui* – *Bathyergus suillus*, *C. hot* – *Cryptomys hottentotus hottentotus*, *C. mah* – *C. h. mahali*, *C. pre* – *C. h. pretoriae*, *F. dam* – *Fukomys damarensis*, *H. gla* – *Heterocephalus glaber* and *G. cap* – *Georychus capensis*.
